# Networks that Link Cytoskeletal Regulators and Diaphragm Proteins Underpin Filtration Function in *Drosophila* Nephrocytes

**DOI:** 10.1101/056911

**Authors:** Simi Muraleedharan, Helen Skaer, Maneesha S. Inamdar

## Abstract

Insect nephrocytes provide a valuable model for kidney disease, as they are structurally and functionally homologous to mammalian kidney podocytes. They possess an exceptional macromolecular assembly, the nephrocyte diaphragm (ND), which serves as a filtration barrier and helps maintain tissue homeostasis by filtering out wastes and toxic products. However, the nephrocyte architecture and elements that maintain the ND are not understood. We show that *Drosophila* nephrocytes have a unique cytoplasmic cluster of F-actin, which is maintained by the microtubule cytoskeleton and Rho-GTPases. A balance of Racl and Cdc42 activity as well as proper microtubule organization is required for positioning the actin cluster. Further, ND proteins Sns and Duf also localize to this cluster and regulate organization of the actin and microtubule cytoskeleton. Perturbation of any of these inter-dependent components impairs nephrocyte ultrafiltration. Thus cytoskeletal components, Rho-GTPases and ND proteins work in concert to maintain the specialized nephrocyte architecture and function.

## Introduction

Chronic kidney disease is a major healthcare problem resulting from gradual loss of ultrafiltration function due to genetic predisposition or disorders such as diabetes and hypertension. The kidney podocyte has a complex morphology that is essential for its role as a glomerular filter. Slit diaphragm (SD) proteins (Neph1, Nephrin) and extracellular matrix-binding transmembrane receptors that make up the filtration diaphragm are coupled to the actin cytoskeleton via integral membrane proteins (Holzman et al., 1999; Horvat et al., 1986; Liu et al., 2003; Sachs et al., 2006). The actin cytoskeleton is a key component that regulates podocyte shape and function. While microtubules (MTs) and intermediate filaments (IFs) provide the major structural support in the podocyte cell body and primary processes (Kobayashi et al., 1998; Kobayashi et al., 2001), dense arrays of actin microfilaments are present in the foot processes (Zhang et al., 2013). In several nephrotic syndromes this differential distribution of cytoskeletal components is lost, leading to loss of SD integrity, foot process effacement and proteinuria (Allison, 2015; Gheissari et al., 2012; Mathieson, 2012; Sanai et al., 2000). RhoGTPase activity orchestrates actin organization and thereby podocyte function (Asanuma et al., 2006; Wang et al., 2012). However, the role of MTs and IFs in regulating the actin cytoskeleton for podocyte function is not well studied.

Recent studies have shown that the *Drosophila* nephrocyte is homologous to the kidney podocyte and has proved to be an excellent molecular-genetic model for elucidating mechanisms that regulate podocyte ultrastructure and function (Tutor et al., 2014; Weavers et al., 2009; Zhang et al., 2013; Zhuang et al., 2009). *Drosophila* nephrocytes, namely pericardial cells (PCs) and garland cells (GCs) flank the cardiac tube and proventriculus respectively (Crossley, 1972; Das et al., 2008c). Nephrocytes carry out ultrafiltration and sequestration of macromolecules, metabolic wastes and toxins from the hemolymph (Das et al., 2008a; Das et al., 2008b). Functional assays of ultrafiltration and lifespan in various *Drosophila* mutants showed that nephrocytes have a filtration diaphragm (nephrocyte diaphragm, ND) that functions in a charge and size selective manner similar to the podocyte SD and is composed of homologous proteins Dumbfounded (Duf; Neph1 homolog) and Sticks and Stones (Sns; Nephrin homolog) (Weavers et al., 2009).

The nephrocyte plasma membrane has extensive invaginations, which resemble the foot processes of podocytes (Crossley, 1972). However, the organization and contribution of various cytoskeletal components in maintaining nephrocyte architecture and function is not known. Understanding these details will make the *Drosophila* nephrocyte a more powerful and valuable model for studying podocyte dysfunction and kidney diseases. Towards this aim we initiated analysis of factors that regulate nephrocyte architecture and function. Here, we show that nephrocyte actin organizes into a unique structure whose position is regulated by the microtubules, Rho-GTPases (Rac1 and Cdc42) and the ND proteins, Sns and Duf. In addition we show that, while microtubules and Sns mutually regulate each other’s expression and localization, Duf does not play a major role in maintaining the microtubule cytoskeleton. Taken together, our results unravel the reciprocal regulation between cytoskeletal components and ND proteins that is essential for size and charge-dependent ultrafiltration.

## Results and Discussion

### Nephrocyte actin is located both cortically and as a central cluster

Since podocyte foot process integrity depends on the maintenance of the actin architecture, we first examined the status of actin in *Drosophila* nephrocytes by staining for F-actin with Phalloidin as well as using an ActinGFP reporter expressed in nephrocytes (*DotGal4>UAS Actin GFP*). GFP expression showed cortical actin at the cell periphery (Fig. 1A, arrowhead). In addition, all nephrocytes showed a medial cluster of densely packed actin located centrally in the cell, adjacent to nuclei (Fig. 1A, white arrow). Phalloidin staining revealed that the actin structures were F actin (Fig 1B). To check whether the actin cluster is tethered to the cell membrane we analyzed transgenic nephrocytes expressing ActinGFP along with membrane localized DsRed (mCD8-DsRed) (*DotGal4>UAS mCD8 Dsred>UAS ActinGFP*). As the nephrocyte plasma membrane invaginates, forming lacunae from which there is rapid endocytosis, mCD8-DsRed is not restricted to the cell periphery (Sup Fig 1A) (Helmstadter et al., 2012). However, the actin clusters showed no co-localization with the mCD8-DsRed indicating that this central actin is not tethered to the membrane. Thus we show for the first time that F actin is present in two compartments of the cell. Cortical actin represents the foot process-like structure and cytoplasmic actin forms a cluster.

**Figure 1.**
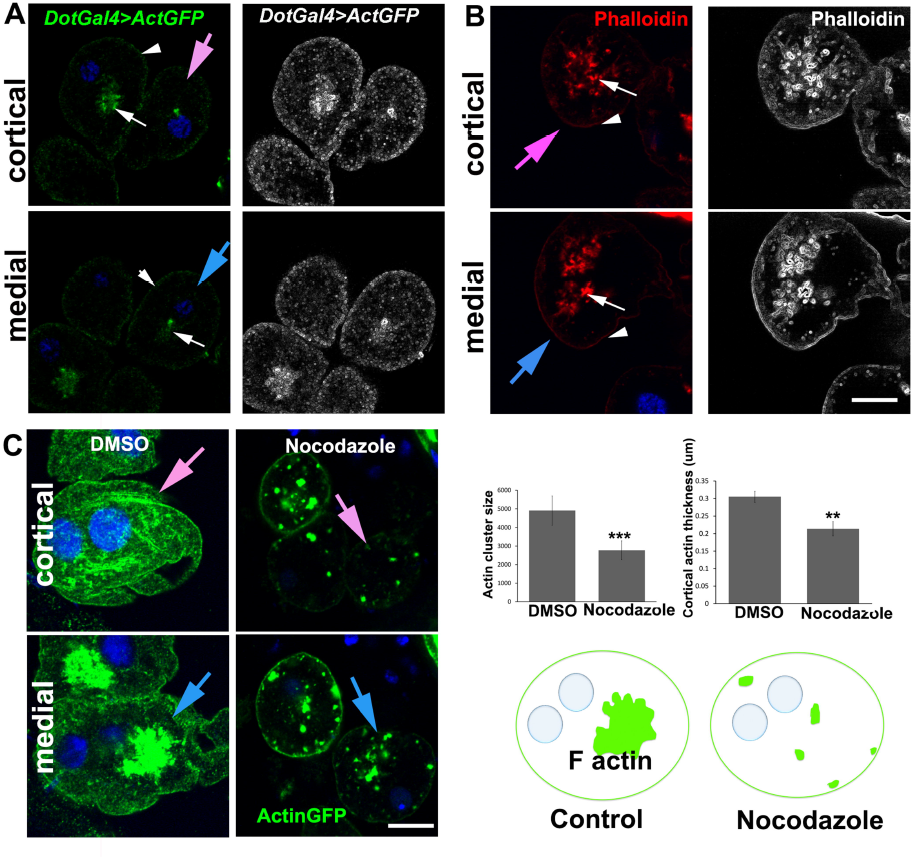
FNephrocytes have cortical actin and a unique actin cluster organization maintained by microtubules. A representative cell at the cortical (pink arrow) or medial (blue arrow) Z slice or projection is indicated. White arrows: actin cluster; arrowheads: cortical actin. (A-B) Actin marked by GFP (green) (*DotGAL4>UAS ActinGFP*) and Phalloidin (red) seen at the cell cortex (arrow head) and as a cytoplasmic cluster (white arrows). (Grey scale images represent the fine edges of actin structures adjusted using ImageJ software) (C) Actin (green) organization after the treatment of nephrocytes with DMSO and Nocodazole. Graph shows the quantification of actin cluster size and cortical actin thickness (see Materials and Methods). Model showing the disintegration of actin cluster in Nocodazole treated cells. Data representative of n>30 from three independent experiments. Error bars depict ± SEM. Scale bars: 10 μm.

### Microtubules regulate actin clusters

Actin and microtubules co-ordinate to maintain and organize cytoplasmic components and organelles (Anesti and Scorrano, 2006; Gurel et al., 2014; Joensuu and Jokitalo, 2015; Kuznetsov et al., 2013). In podocytes, microtubules are present in and maintain the primary foot processes (Kriz et al., 1995) (Saleem et al., 2002). In nephrocytes, both GFP (*DotGal4>UAS Tubulin* GFP) and immunolabelling shows that Tubulin is largely punctate but in some regions appears filamentous (Sup Fig 1B). Co-staining of microtubules and actin showed that although the two co-localized at the cortex, actin clusters are devoid of microtubules (Sup Fig 1C). In order to assess the effect of microtubule disruption in nephrocytes, we standardized Nocodazole treatment assay in nephrocytes and found that disruption of microtubules caused disintegration of actin clusters and decreased cortical actin (Fig. 1C), suggesting that the maintenance of actin requires intact microtubules. This indicates that microtubules regulate actin organization in nephrocytes.

### A balance of Racl and Cdc42 activity is required for actin organization

Balanced activity of the Rho-GTPases RhoA, Rac1 and Cdc42 is essential for maintaining podocyte architecture and function (Mouawad et al., 2013).

To test the role of Rho-GTPases in nephrocytes, we impeded their activity by expressing dominant negative (DN) forms of Rac1 (*DotGal4>UAS-RacN17*), Cdc42 (*DotGal4>UAS-Cdc42N17*) and Rho A (*DotGal4>UAS-RhoN19*) and assayed for the effect on cortical actin and actin clusters (Fig. 2A). Quantitative analysis was done based on measuring the size of actin cluster and thickness of cortical actin. Lack of Rac1 activity obliterated actin clusters, whereas Cdc42-DN nephrocytes showed several smaller dispersed clusters with a significant increase in actinGFP intensity as calculated by the size of each actin cluster. DN RhoA did not affect actin clusters. Thus Rac1 is essential for the formation and/or maintenance of actin clusters and possibly actin stability. Cdc42 has a negative effect on actin stability and could limit the size and distribution of actin clusters. Upon Rac1 or Cdc42 inactivation, we also found that cortical actin thickness is noticeably affected with decreased thickness in Racl DN and an increase in the Cdc42DN nephrocytes. Hence, in nephrocytes, a balance of activity of Racl and Cdc42 activity is required for the unique actin organization (Fig. 2B). Our study shows the importance of Rho-GTPases in cytoskeletal regulation of *Drosophila* nephrocytes and lays the foundation for identifying specific regulators.

**Figure 2.**
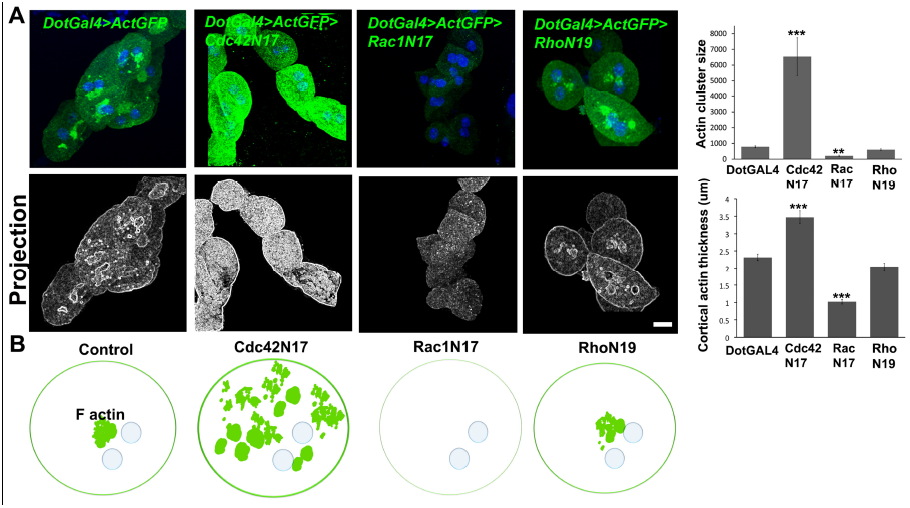
Racl and Cdc42 regulate actin cluster organization. (A) ActinGFP (green) in control and Cdc42N17, RaclN17 and RhoN19 DN nephrocytes as indicated. Graphs represent the actin cluster size and cortical actin thickness. Actin cluster size and cortical actin thickness were increased in the Cdc42 dominant negative nephrocytes and decreased in Racl dominant negative cells. (B) Models representing actin organization in control, Cdc42N17, RaclN17 and RhoN19 expressing nephrocytes. Cdc42N17 nephrocytes show increased cortical actin thickness and actin cluster size (green), whereas the RaclN17 nephrocytes show thinner cortical actin and nearly an absence of the actin cluster and RhoN19 nephrocyte shows no change. Data representative of n>30 from three independent experiments. Error bars depict ± SEM. Scale bars: 10μm.

### Nephrocyte diaphragm proteins are essential for nephrocyte actin organization

Podocyte SD proteins Nephrin and NEPH1 are transmembrane proteins that signal to activate a myriad of pathways that regulate actin organization in foot processes. Nephrocyte diaphragms have homologous proteins, namely Kirre/Dumbfounded (Duf) and Roughest (Rst) proteins (homologous to NEPH1); Hibris (Hbs); Sticks and stones (Sns) (homologous to NPHS1) (Weavers et al., 2009; Zhuang et al., 2009). However their interaction with and ability to regulate the organization of actin have not been reported. Hence, we tested the effect of perturbing representative ND proteins Sns and Duf, on actin organization. Protein localization and ActinGFP expression showed that Sns and actin co-localize at the cell cortex and cluster whereas Duf co-localizes with cortical actin but co-habits with the actin clusters (Fig 3A). This indicates that there is a pool of intracellular diaphragm proteins, whose distribution overlaps with actin clusters. Depletion of the ND proteins (*DotGal4>UAS SnsRNAi; DotGal4>UAS DufRNAi*) showed that lack of Sns caused dramatic disorganization of actin so that clusters were lost and cortical actin thickness was strongly reduced (Fig 3B). The reduction in number of clusters and cortical actin on Duf depletion was moderate but significant. Together these data suggest that actin organization is dependent on the ND proteins.

**Figure 3.**
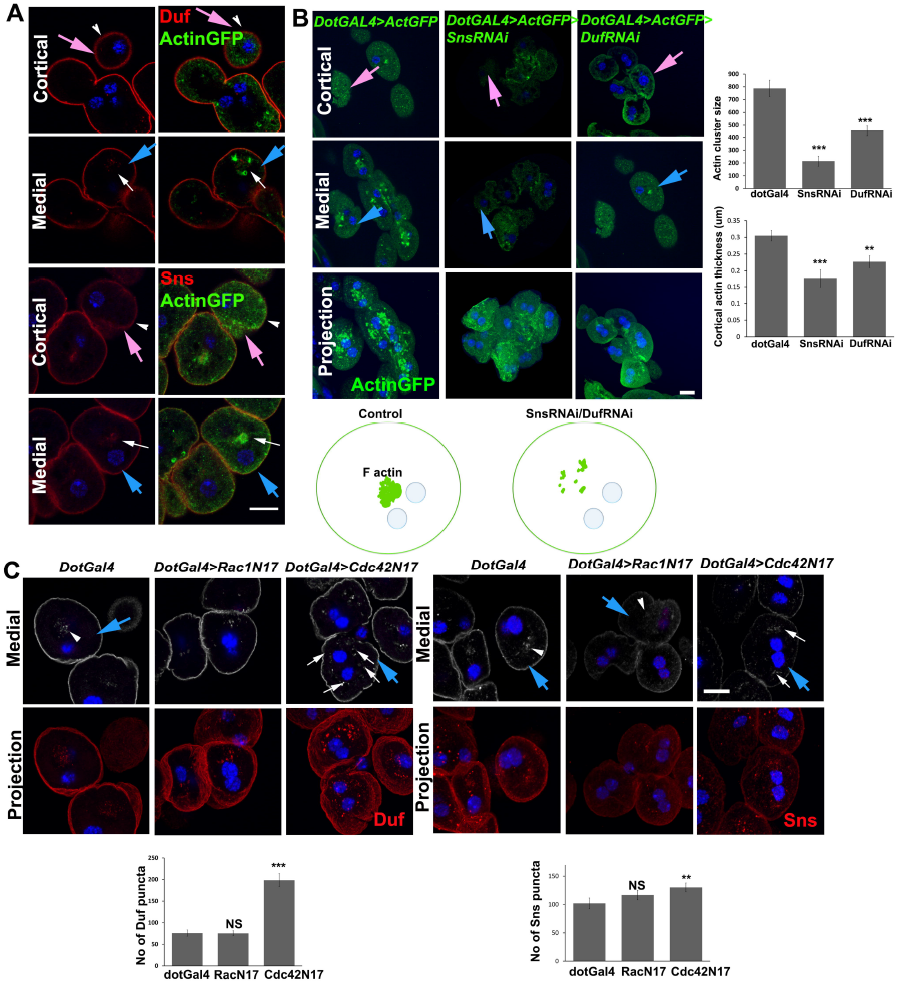
Actin organization and nephrocyte protein localization are interdependent. A representative cell at the cortical (pink arrow) or medial (blue arrow) Z slice or projection is indicated. (A) Actin (green) and Duf or Sns (red) co-staining in garland cells. ND proteins (red) and actin associate at the cortex (arrow heads) as well as at the cluster (arrows). (B) Actin (green) organization in Sns RNAi (*DotGAL4>UASSnsRNAi*) and Duf RNAi (*DotGAL4>UASDufRNAi*) nephrocytes. Graph represents quantification of actin cluster size in control and ND-RNAi garland cells. Models showing that ND proteins localize near the actin cluster, which is perturbed in ND mutants. (C) Duf and Sns localization pattern in control, RacN17 and Cdc42N17 expressing garland cells. Duf and Sns cytoplasmic puncta are increased (arrows) in the Cdc42N17 and dispersed (arrowhead) in RacN17 nephrocytes. Graphs represent quantification of Duf and Sns puncta. Data representative of n>30 from three independent experiments. Error bars depict ± SEM. Scale bars: 10μm.

### Racl and Cdc42 regulate ND proteins

Balanced activity of all the Rho-GTPases is essential for foot process organization as well as slit diaphragm arrangement in podocytes (Attias et al., 2010; Scott et al., 2012). Analysis of ND proteins Sns and Duf in DN Rho-GTPase-expressing nephrocytes showed that Racl inactivation led to dispersed Sns and Duf punctae with no significant change in number. However, Cdc42-DN caused significantly more Sns and Duf puncta (Fig. 3C). This correlates well with the increased actin cluster phenotype seen upon Cdc42 inactivation. Thus proper arrangement of actin clusters is essential for the normal cytoplasmic localization of ND proteins and actin organization and ND protein localization are interdependent.

### Sns and microtubules regulate each other in nephrocytes

Our analysis shows that both tubulin and ND proteins regulate actin organization in nephrocytes. Hence we tested whether ND proteins and microtubules affect each other’s organization. Nephrocytes depleted of Sns but not of Duf, showed reduced tubulin staining (Fig 4A). Conversely, when microtubules were disrupted using Nocodazole, there was a decrease in the cortical as well as the cytoplasmic Sns while Duf localization remained comparable to the control (Fig 4B). This indicates that Sns and microtubules are interdependent for their organization suggesting specificity in the interaction.

**Figure 4.**
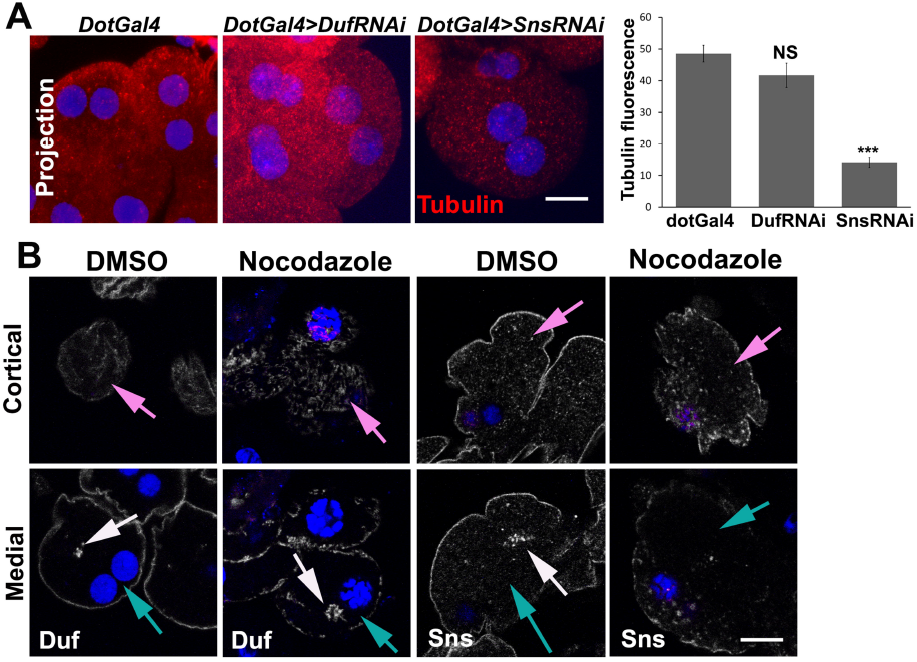
Sns and microtubules regulate each other in nephrocytes. A representative cell at the cortical (pink arrow) or medial (blue arrow) Z slice or projection is indicated. (A) Tubulin staining (red) in Duf RNAi (*DotGAL4>UASDufRNAi*) and Sns RNAi (*DotGAL4>UASSnsRNAi*) nephrocytes. Graph represents quantification of Tubulin fluorescent intensity. Microtubules are drastically reduced in the SnsRNAi but not in DufRNAi nephrocytes. (B) Duf and Sns (white) staining in DMSO and Nocodazole-treated garland cells. Sns localization is disrupted in Nocodazole treated cells while Duf distribution is moderately affected (white arrows). Data representative of n≥30 from three independent experiments. Error bars depict ± SEM. Scale bars: 10μm.

### Actin cluster regulates endoplasmic reticulum morphology in nephrocytes

Cortical actin represents the nephrocyte foot processes and it is directly linked to filtration function. In order to understand the significance of the actin cluster, we analyzed the localization pattern of actin cluster with cellular organelles and particularly found that actin cluster was placed adjacent to endoplasmic reticulum (ER) structure (Fig 5A). Many reports suggest that cytoskeleton regulates ER to maintain the ER tubules and sheets in different cell types (Joensuu et al., 2015). To understand the regulation between ER and the specific actin cluster in nephrocytes, we observed ER organization using an ER lumen marker, Boca, in control and Racl, Cdc42, Sns and Duf mutants (*DotGal4>UAS-RacN17; DotGal4>UAS SnsRNAi*; *DotGal4>UAS DufRNAi; DotGal4>UAS-Cdc42N17; DotGal4>UAS-RhoN19*). ER intensity was drastically increased in the Rac1 DN mutant and moderately increased in Cdc42 DN mutant. Interestingly, Duf RNAi did not show significant change but Sns RNAi showed drastic reduction of ER (Fig. 5B). These results suggest that actin cluster plays a role in the maintenance of ER morphology in nephrocytes.

**Figure 5.**
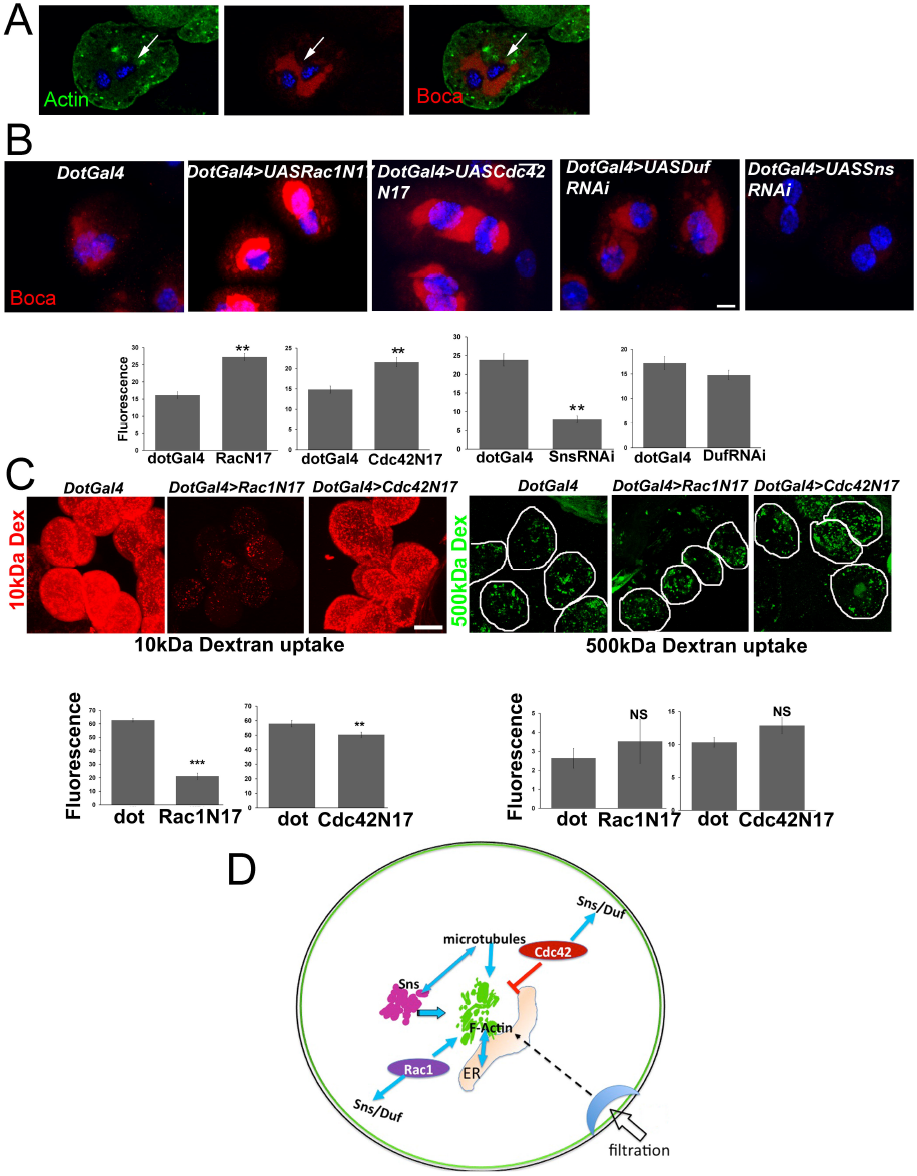
FActin cluster maintains endoplasmic reticulum organization in nephrocytes. (A)Actin (green) and ER lumen marker, Boca (red) co-staining in nephrocytes. Actin cluster lies adjacent to the ER. (B) ER intensity was drastically increased in the Racl DN and moderately increased in the Cdc42 DN mutant compared to the control. But SnsRNAi showed reduced ER and Duf RNAi did not affect the ER. Graphs showing the Boca fluorescence intensity in the control and mutants. (C) Uptake of 10 kDa Texas Red Dextran (red) or 500 kDa FITC Dextran (green) in control (*DotGal4*), RacN17 and Cdc42N17 expressing nephrocytes. 10 kDa Dextran uptake was reduced drastically in RaclN17 nephrocytes and slightly in Cdc42N17 nephrocytes. 500 kDa Dextran uptake was not significantly changed. Graphs represent the quantification of total fluorescence intensity. (D) Model summarizing regulation of nephrocyte actin by microtubules, Rho-GTPases and ND proteins for proper ER organization. Data representative of n>30 from three independent experiments. Error bars depict ± SEM. Scale bars: 10μm.

### Rho-GTPases aid in ultrafiltration

Disruption of microtubules, ND protein or Rac1 or Cdc42 perturbs actin structure. In podocytes, perturbed RhoGTPase activity (and hence actin organization) leads to massive proteinuria and foot process effacement. Hence we tested the effect of disrupting actin organization on nephrocyte function by assaying for ultrafiltration in dominant negative Racl or Cdc42 nephrocytes. Interestingly inactive Rac1 and Cdc42 did not affect the uptake of larger (500 kDa) Dextran molecules but uptake of 10 kDa Dextran was compromised (Fig 5C). Thus Rac1 and Cdc42 regulate size-based ultrafiltration in nephrocytes. This indicates that actin organization as well as ND protein organization is important for nephrocyte function. Cortical actin is likely to be important for the plasma membrane folding and the maintenance of lacunae and SDs. Thus both cortical and central clusters of F-actin may have an essential role in maintaining nephrocyte ultrastructure for function.

This study highlights the role of intracellular organelles and nephrocyte diaphragm proteins in the regulation of the nephrocyte cytoskeleton for the maintenance of its unique structure. We provide previously undocumented evidence for the distribution of actin at the cell cortex and in the form of cytoplasmic clusters in nephrocytes and show that the tight regulation of this cytoskeleton structure is required for nephrocyte filtration function. The more complex mouse podocyte architecture with secondary and tertiary foot processes is especially vulnerable to cytoskeletal defects and depended on RhoA, Rac1 and Cdc42. In contrast, we show that nephrocyte actin is maintained by the balanced activity of Rac1 and Cdc42.

In nephrocytes Sns and Duf co-express in the same cell and are essential for diaphragm and foot process formation (Weavers et al., 2009). However, in *Drosophila* eye and muscle cells, Sns and Duf are expressed in two different cell types and function in a complementary fashion. Our study additionally shows the significance of cytoplasmic Sns and Duf in regulating actin organization, which could be direct or a result of altered cortical actin when ND proteins are lost or reduced. Further, Sns has a greater role in nephrocyte architecture and function. Cultured kidney podocytes also show punctate, cytoplasmic localization of SD proteins, which partially overlap with the actin stress fibers (Saleem et al., 2002). Thus such a cytoplasmic correlation is conserved but the mechanism and function is not known yet.

The intricate connection between endoplasmic reticulum and cytoskeleton is established in different cell types. Actin and microtubules associate with the ER in a specialized manner forming a network to regulate ER structure. In this study, we observe the relationship between ER and nephrocyte cytoskeleton. The actin cluster lies adjacent to the ER and such a localization is necessary for proper ER organization. When the actin cluster is disrupted, the ER expands. Similarly, Sns plays a major role in ER maintenance possibly through microtubules and actin. Thus multiple structural components crosstalk to maintain the complex and specialized architecture of the nephrocyte, thereby regulating efficient filtration function. This study is relevant to kidney podocytes where slit diaphragm proteins and the cytoskeleton play significant roles in the filtration function, but the crosstalk between cytoskeleton and ER is largely unexplored area.

In summary, our findings elucidate the conserved molecular interactions in nephrocytes that are essential for ultrafiltration function and increase its relevance as a model for kidney disease. This report provides fundamental information required to probe further the cross talk between cytoskeletal elements and ND proteins, which could aid analysis of podocyte biology and kidney disease.

## Methods and materials

***Drosophila* stocks used:** Canton-S, *UAS Actin GFP (III*) (Bloom 9257); *UAS TubulinGFP* (Bloom 7373); UAS *RacN17* (Bloom 6292); *UAS Cdc42N17* (Bloom 6288); *UAS RhoA N19* (Bloom 7328); *UAS snsRNAi*; *UAS DufRNAi*.

**Immunostaining and immunofluorescence analysis:** Dissected nephrocytes were immunostained with desired primary and secondary antibodies as described before (Das et al., 2008c). Antibodies used: Chick-and rabbit-anti-GFP (Invitrogen, 1:500); mouse antiTubulin (DSHB, 1:100); Rat anti-Duf (Eyal Schejter, Weizmann Institute of Science, Israel); Rabbit anti-Sns (Susan Abmayr, SIMR, USA), Guinea pig anti Boca (1:100) (Richard Mann, Columbia University, USA). Phalloidin-Alexa568 (Invitrogen, 1:100) was used to detect actin. Imaging was on Zeiss LSM-Meta510 or LSM780.

**Nocodazole treatment assay:** Dissected garland cells (N=10) were incubated in S2 medium containing various concentrations of Nocodazole or DMSO (Sigma Aldrich) for 6 hours at room temperature, fixed in 4% paraformaldehyde and immunostained. 1 mM Nocodazole was selected for experiments based on anti-tubulin staining.

**Quantification and statistical analysis:** n>/=30 for each genotype in each experiment. Actin cluster quantitation was done by calculating the particle size, number and area using Image J software. Average values were plotted using Microsoft Excel. Statistical significance was calculated using single factor ANOVA (analyses of variance) with Microsoft Excel (P<0.01 was considered significant).

**Ultrafiltration assay:** Refer to (Weavers et al., 2009).

**Co-localization index calculation:** Single optical slices were analyzed using the Histo option in the Zeiss examiner LSM software and the co-localization graphs were plotted.

## Acknowledgments

We thank *Drosophila* community for fly stocks and antibodies, Imaging Facilities at JNCASR and NCBS for access and our laboratory members for fruitful discussion. Funding was from grant no. 094879/B/10/Z from The Wellcome Trust, UK to HS and MSI and intramural funds from JNCASR to MSI.

## Author contributions

S.M., M.S.I. designed research; S.M. performed experiments; S.M, M.S.I, H.S analyzed data; S.M, M.S.I, H.S wrote the paper.

## Supplementary figures

**Sup Figure 1.**
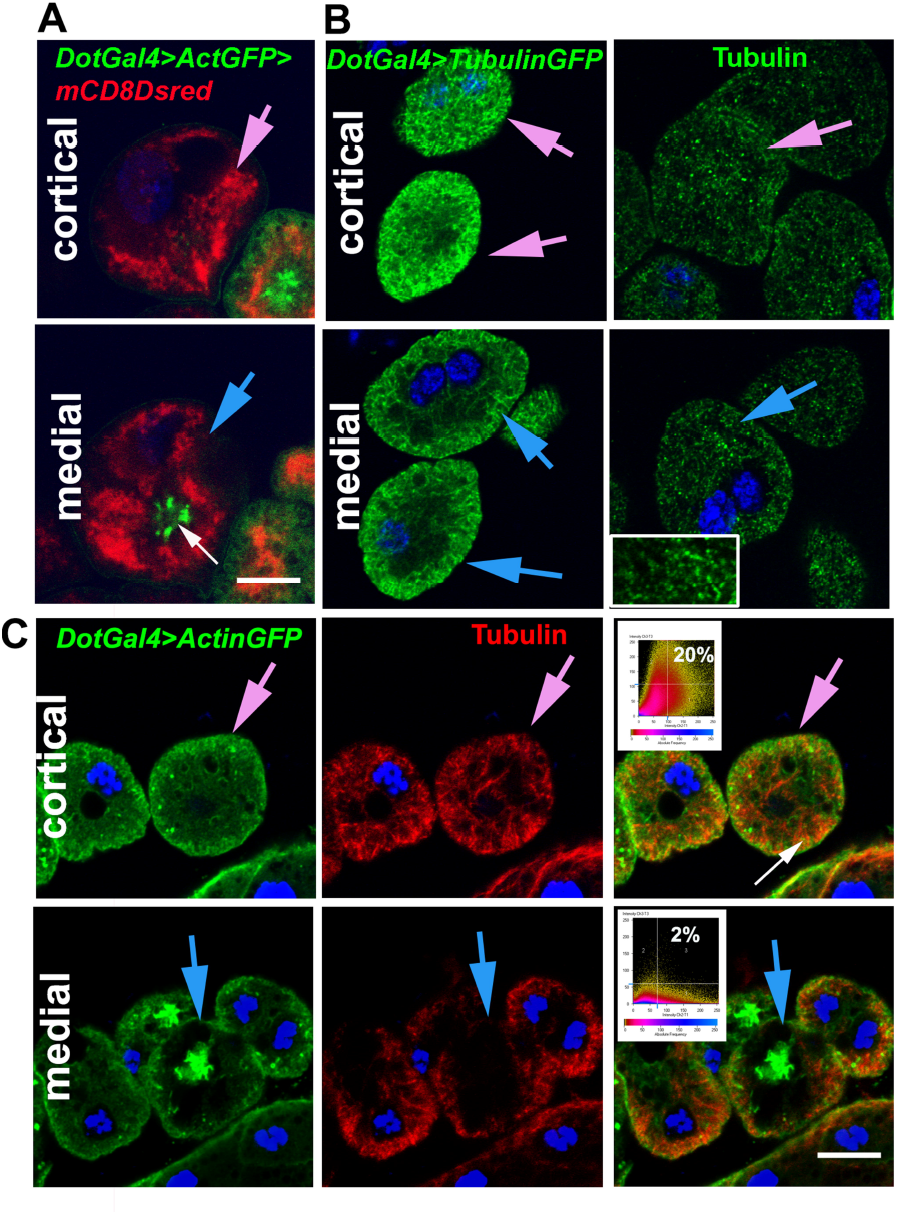
Microtubules do not co-localize with actin cluster. (A) Membrane (red) and actin (green) in nephrocytes expressing *DotGAL4>mCD8 dsred>ActinGFP* showing that actin cluster do not tether to the cell membrane.(B) Tubulin staining (green) of nephrocytes in DotGal4>TubulinGFP and CantonS (inset showing the partial punctate appearance of microtubules in nephrocytes) (C) Actin (green) and Tubulin (red) partial co-localization (arrows) at the cortex but not at the actin cluster (Inset: co-localization plots). Data representative of n>30 from three independent experiments. Scale bars: 10μm.

